# Annotation of Spatially Resolved Single-cell Data with STELLAR

**DOI:** 10.1101/2021.11.24.469947

**Authors:** Maria Brbić, Kaidi Cao, John W. Hickey, Yuqi Tan, Michael P. Snyder, Garry P. Nolan, Jure Leskovec

## Abstract

Accurate cell type annotation from spatially resolved single cells is crucial to understand functional spatial biology that is the basis of tissue organization. However, current computational methods for annotating spatially resolved single-cell data are typically based on techniques established for dissociated single-cell technologies and thus do not take spatial organization into account. Here we present STELLAR, a geometric deep learning method for cell type discovery and identification in spatially resolved single-cell datasets. STELLAR automatically assigns cells to cell types present in the annotated reference dataset as well as discovers novel cell types and cell states. STELLAR transfers annotations across different dissection regions, different tissues, and different donors, and learns cell representations that capture higher-order tissue structures. We successfully applied STELLAR to CODEX multiplexed fluorescent microscopy data and multiplexed RNA imaging datasets. Within the Human BioMolecular Atlas Program, STELLAR has annotated 2.6 million spatially resolved single cells with dramatic time savings.

## Introduction

Development of spatial protein and RNA imaging technologies has opened new opportunities for understanding location-dependent properties of cells and molecules [1–4]. The power to capture spatial organization of cells within tissue plays an essential role in understanding cellular function and in studying complex intercellular mechanisms. To increase our knowledge of cells in healthy and diseased tissues, large tissue-mapping consortia efforts such as the Human BioMolecular Atlas Program (HuBMAP) [5], Human Tumor Atlas Network (HTAN) [6], and Human Cell Atlas (HCA) [7] have been generating comprehensive cell atlas datasets. These consortia efforts necessitate computational methods that can assist with robust characterizations of cells and guide our understanding of functional spatial biology. While computational tools for dissociated single-cell technologies have been used to characterize cells in spatial datasets [4, 8–10], these tools ignore the spatial information crucial for more accurate annotation of spatial datasets.

Recently, methods that take into account spatial organization of cells have been developed for annotating spatial transcriptomics data [11–15]. However, these methods can not automatically assign cell type labels to cells and thus require human reannotation once clusters are identified. While CELESTA tool has successfully avoided post hoc cluster reannotation for multiplexed in situ images [16], it is reliant upon the prior human supervision provided in the form of the set of marker genes of expected cell types which is often challenging to define for a new dataset. As new spatial datasets are being generated [17–21], there is a necessity for computational methods that simultaneously leverage molecular features and additional spatial context of cells while at the same time minimize manual human annotation effort.

Here, we present STELLAR (SpaTial cELl LeARning), a geometric deep learning tool for cell-type discovery and identification in spatially resolved single-cell datasets. Given annotated spatially resolved single-cell dataset with cells labeled according to their cell types (reference dataset), STELLAR learns spatial and molecular signatures that define cell types. Using the reference dataset, STELLAR then transfers the annotations to a completely unannotated spatially resolved single-cell dataset with unknown cell types. The reference and unannotated datasets can belong to different dissection regions, different donors, or different tissue types.

STELLAR has two unique properties. First, using graph convolutional neural networks [22, 23], STELLAR learns latent low-dimensional cell representations that jointly capture spatial and molecular similarities of cells. In this way, cells that are spatially close together and that have similar levels of gene or protein expression are embedded close to each other. Indeed, we show that cell embeddings learnt by STELLAR reveal higher-order tissue structures. Second, in a new completely unannotated dataset, STELLAR automatically assigns cells to cell types included in the reference set and also identifies cells with unique properties as belonging to a novel type that is not part of the reference set. Thus, STELLAR has an ability to assign cells to one of the cell types seen in the reference set, or discover a novel cell type for previously uncharacterized cell types. From a practical standpoint this ability is essential for leveraging reference sets generated in a disease-free state or in different tissues that may not have all the cell types represented in the target tissue but that may include most established cell types.

As a result of these unique properties, STELLAR solves major limitations with current annotation tools. STELLAR can be applied to both multiplexed protein and multiplexed RNA imaging datasets. In particular, we show the effectiveness of STELLAR on annotating CODEX multiplexed fluorescent microscopy datasets and MERFISH spatial transcriptomics datasets. Encouraged by the results, we used STELLAR to annotate HuBMAP consortium data: so far STELLAR has annotated 2.6 million cells labeled with 54 protein markers from 8 different regions of the human intestine data and across 8 different donors [17]. On these datasets, STELLAR has saved hundreds of hours of manual work needed to assign cells to cell types.

## Results

### Overview of STELLAR

STELLAR takes as input *(i)* The reference dataset of annotated spatially resolved single cell-dataset with cells assigned to cell types, and *(ii)* a completely unannotated spatially resolved single-cell dataset with cell types that are unknown (Fig. 1). STELLAR then assigns unannotated cells to cell types in the reference dataset, while for cells that do not fit into the reference cell types, it identifies individual novel cell types and assigns cells to them. STELLAR learns low-dimensional cell representations (embeddings) that reflect molecular features of cells as well as their spatial organization and neighborhood. The cell embeddings are learnt using graph convolutional neural networks (GCNs) [22, 23] – neural networks that enable learning over arbitrary graph structures by encoding the structure of the local graph neighborhood into a dense vector embedding. In STELLAR, we construct a graph based on the spatial proximity of cells and use molecular features of cells as node features (Methods).

**Figure 1:**
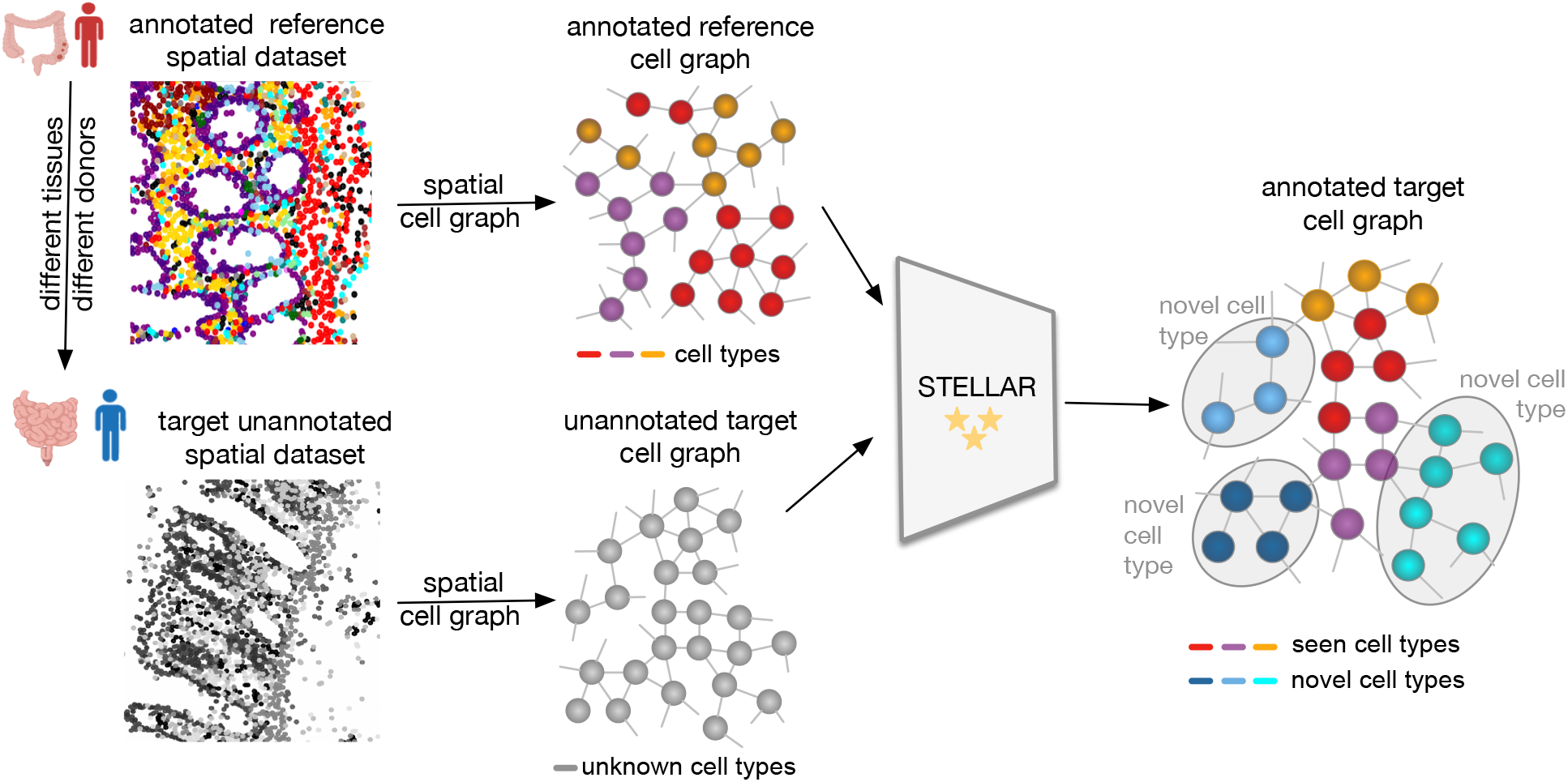
STELLAR is a geometric deep learning framework for annotating spatially resolved single-cell datasets. Given a reference spatially resolved single-cell dataset in which cells are annotated according to their cell types, STELLAR assigns cells in an unannotated spatial single-cell dataset to cell types included in the reference dataset or discovers novel cell types as a group of cells with unique properties not present in the reference dataset. STELLAR uses a graph convolutional encoder to learn low-dimensional cell embeddings that capture cell topology as well as their molecular profiles. Reference and unannotated datasets can originate from different sources as STELLAR transfers information across different tissues and different donors.

Given cell embeddings, STELLAR has two capabilities: (1) it assigns cells in the unannotated dataset to one of the cell types previously seen in the reference dataset; And (2), for cells that do not fit the existing known/labeled cell types STELLAR identifies novel cell types and assigns cells to them. STELLAR achieves this using an objective function that consists of two main components (Extended Data Fig. 1). In the first component, STELLAR gradually learns to separate cell types from the reference set by controlling intra-class variance of previously seen cell types using the adaptive margin mechanism [24]. In the second component, STELLAR discovers novel classes by generating auxiliary labels for the unannotated data which are used to guide the training. The auxiliary labels are generated based on the nearest neighbors of cells in the embedding space (Methods).

### STELLAR identifies existing and discovers novel cell types

To demonstrate STELLAR’s ability to assign cells to one of the cell types seen in the reference set and discover a novel cell type, we applied STELLAR on CODEX multiplexed imaging data [10]. We used data from human tonsil as the reference set (Fig. 2a and Extended Data Fig. 2) and tissue from a patient with Barrett’s esophagus (BE) as the unannotated target tissue (Fig. 2b). Both imaging datasets underwent image processing and single cell segmentation. Cell type labels were manually curated and assigned by iterative unsupervised clustering of the single cell data, analyzing expressions of protein markers, and validating cluster accuracy with visualization in the tissue. Tonsils are often used for testing antibody panels as they contain high numbers of immune cell markers. The BE tissue also contains immune cells, but additionally contains differentiated epithelial cells that tonsils do not. Indeed, three subtypes of epithelial cells appear only in the BE dataset, while B cells appear only in tonsil data. Moreover, the two datasets have different distributions of cell types. For example, smooth muscle cells are major cluster in BE, but form the smallest group in the tonsil dataset (Extended Data Fig. 3).

**Figure 2 (preceding page):**
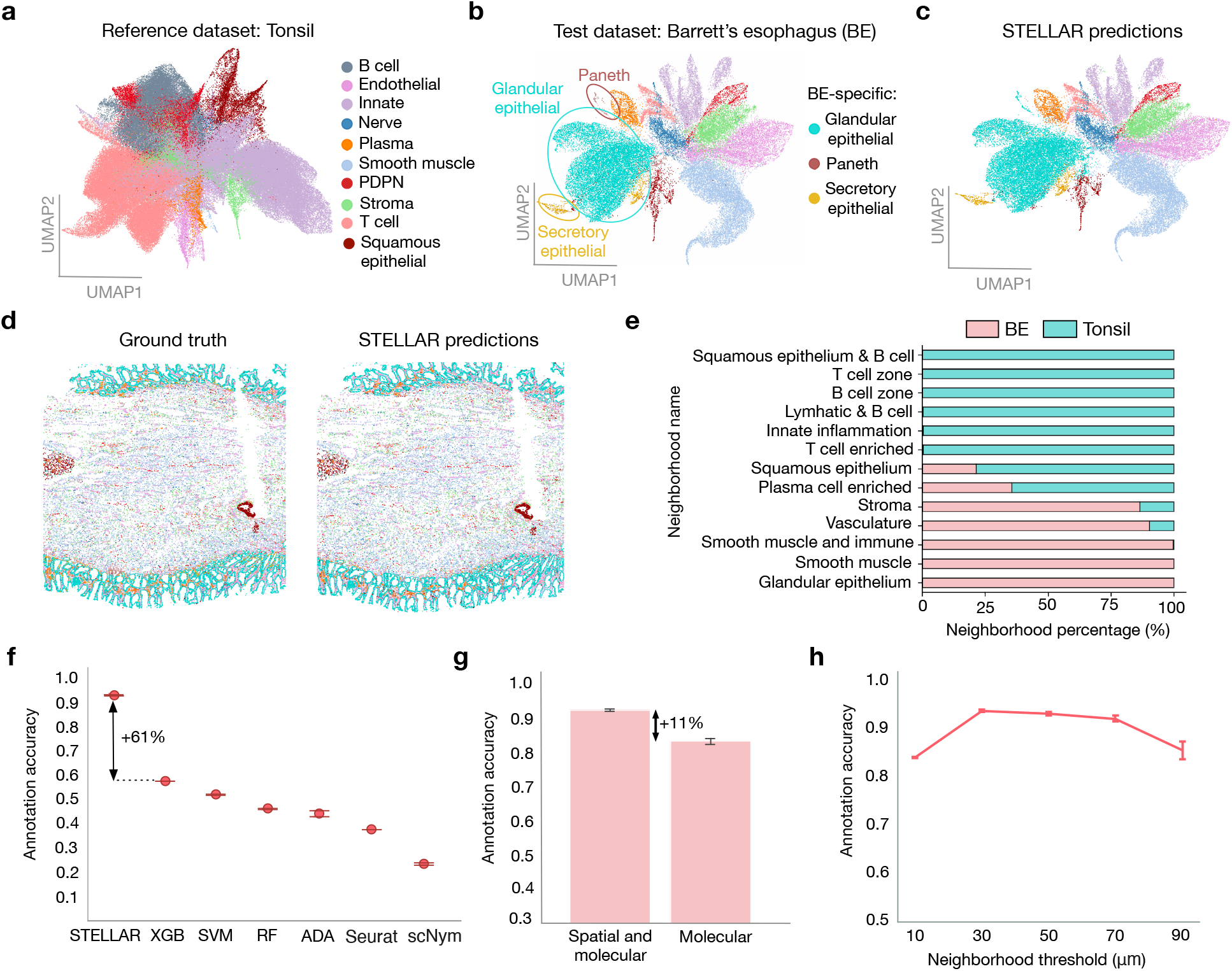
STELLAR accurately identifies cell types from the reference set and discovers novel cell types that have never been characterized in the reference set. **(a)** UMAP visualization of the healthy tonsil data used as the reference dataset. Colors denote ground truth cell types annotations used to train STELLAR. **(b, c)** UMAP visualization of Barrett’s esophagus (BE) data used as the test set. Three subtypes of epithelial cells are not found in the tonsil reference data. Colors denote **(b)** ground-truth cell-type annotations and **(c)** STELLAR predictions. **(d)** CODEX image of BE in spatial coordinates colored according to ground-truth annotations (left) and STELLAR predictions (right). **(e)** Different neighborhood composition between reference dataset from tonsil tissue and target unannotated dataset from BE. Neighborhoods are determined by cell composition enrichment in nearest neighbor vectors for each cell. **(f)** Accuracy of cell-type assignments in the BE dataset by STELLAR compared to alternative approaches. Position of scatter plot points is computed as a mean accuracy score across five runs of each method. Error bars are from standard deviation. **(g)** Accuracy of cell-type assignments in BE dataset by STELLAR and when modifying STELLAR to rely only on the molecular information in cell type assignment (Methods). Height of each bar is computed as a mean accuracy score across five runs of each method. Error bars are from standard deviation. **(h)** The effect of neighborhood threshold on the ‘s performance. All cells with distance less than the threshold are considered as neighbors. Position of scatter plot points is computed as a mean accuracy score across five runs of each method. Error bars are from standard deviation.

Despite these major differences, we found that STELLAR accurately assigned cell types to 93% of cells in the BE data, discovering also BE-specific subtypes of epithelial cells as a newly discovered cell type and correctly differentiating between glandular epithelial and secretory epithelial novel cell types (Fig. 2c). Novel cluster of paneth cells was correctly recognized as a novel class by STELLAR but assigned to glandular epithelial cells. The reason is that paneth cells are very rare in the BE dataset and form the smallest class of only 0.6% of the total cell number in the BE dataset. Mapping cell type annotations back to spatial coordinates, STELLAR predictions also show agreement with ground truth annotations and do not reflect issues recognizing cell types in certain areas of the tissue (Fig. 2d). The disagreement with ground truth annotations mostly comes from mixing major cell types, in particular endothelial with stroma cells and endothelial with smooth muscle cells (Supplementary Fig. 1). These are all stromal cells and they likely share Vimentin, lacking CD45 or Cytokeratin. They are also very often spatially located next to each other – endothelial cells will often be next to smooth muscle cells and vice versa. Furthermore, endothelial cells are elongated and the problem may be caused by current segmentation software tools that co-segment them with other cell type markers causing STELLAR to assign them to other cell types.

One of the reasons we applied STELLAR to these two datasets was also to test the flexibility of the algorithm when cells may be found in different cellular environments. This is important for cross-tissue applications, but also in cases of applying STELLAR across disease states like cancer where cells, normally restricted to one area, invade other areas of the tissue. Since we observed a high accuracy in cell type label transfer, we asked whether labels that STELLAR generates result in unique cellular microenvironments or neighborhoods. This would mean that STELLAR is able to generalize even when the same cell types are present in a different microenvironment. To explore this, we used a method for cellular neighborhood analysis [25] with the ground truth labels of the tonsil dataset and the STELLAR’s predictions for the BE dataset. This method determines the composition of the nearest neighbors for each cell. Then these composition vectors for each cell are clustered into similarly composed cellular neighborhoods (Methods). Interestingly, this analysis revealed both shared and unique multicellular neighborhoods (Fig. 2e, Extended Data Fig. 4). For example, glandular epithelium and smooth muscle are exclusively found within BE tissue, while B cell zone, T cell zone, and squamous epithelium/B cell neighborhoods are only found in the tonsil dataset. Furthermore, even many neighborhoods shared between the datasets have little overlap in total percent of cells indicating the structural distinctness of the two tissues. Thus, STELLAR can generalize to different spatial organizations while accurately transferring cell type labels.

### STELLAR outperforms other methods

On the BE dataset, we compared STELLAR to four other baselines including XGBoost [26], Support Vector Machine (SVM) [27], Random Forest [28], AdaBoost [29], as well as single-cell annotation baselines Seurat V4 [30] and scNym [31]. We trained all baselines on the tonsil dataset, and evaluated them on the BE dataset. STELLAR substantially outperformed the best alternative method, achieving 61% higher accuracy (Fig. 2f and Supplementary Fig. 2). Single-cell baselines Seurat V4 and scNym achieved lower performance than other baselines so we additionally compared to another single-cell baseline scANVI [32]. However, the performance of scANVI was even lower, so we created a leaky setting for scANVI in which we used a fraction of BE annotations for model training. In such a scenario, the performance of scANVI substantially improved to 0.76 indicating that the performance drops are caused by differences between tonsil and BE datasets (Supplementary Methods and Extended Data Fig. 5a). Even when using BE annotations to annotate BE dataset, scANVI still did not outperform STELLAR. STELLAR was also the best performing method when evaluating performance using other evaluation metrics, such as F1-score, precision, and recall (Extended Data Fig. 5 b-d). To directly measure performance on cell types that are only present in the reference dataset, *i.e.,* shared between tonsil and BE datasets, we removed novel cell types. In this scenario we still find that STELLAR outperformed other alternative methods by 6% (Supplementary Fig. 3).

We further systematically tested robustness of STELLAR to: *(i)* different data normalization strategies, *(ii)* artificially introduced noisy annotations, and *(iii)* different number of withheld marker proteins. First, we find that STELLAR achieves best performance on z-score normalized CODEX data which agrees with previous studies comparing different combinations of normalization techniques and unsupervised clustering [33] (Extended Data Fig. 6a). STELLAR outperforms alternative methods even with different normalization strategies including when evaluated on unnormalized data, achieving 87% accuracy on unnormalized data. We additionally evaluated the effect of the noisy annotations on the performance. Specifically, we misannotated proportion of cells by randomly assigning wrong annotations to cells in the reference tonsil dataset. With 5% of wrong annotations, STELLAR achieves only 0.7% lower performance compared to performance in which all annotations are correct (Extended Data Fig. 6b). This means that STELLAR is robust even when proportion of cells are misannotated. Finally, STELLAR is ran on the shared subsets of the genes/proteins between annotated reference dataset and unannotated target dataset. STELLAR is able to accurately transfer cell type labels even when leaving out measured marker proteins (Extended Data Fig. 6c). While not as critical for transcriptomic datasets, this may be important for transferring cell type labels in multiplexed imaging where there are limited numbers of antibody markers that can be chosen and antibody panels may not completely overlap.

### Importance of capturing spatial organization of cells

Using graph convolutional neural networks, STELLAR leverages both spatial organization of cells as well as their molecular expressions. To directly measure the benefits of including spatial information, we designed a baseline in which we used STELLAR’s objective function, but we replaced graph convolutional encoder with a fully connected neural (FCN) network layer disabling the usage of spatial information. When comparing FCN baseline to STELLAR, we find that capturing spatial organizations of the cells brings 11% improvement in cell type annotation task compared to relying solely on the molecular information (Fig. 2g).

We next sought to analyze the optimal graph structure for the task. In STELLAR, we construct the graph by considering as neighbors all the cells whose distance is less than the preset threshold which we initially set to 50 *μm*. We systematically changed the value of the threshold and analyzed the effect on the STELLAR performance on the CODEX tonsil/BE dataset. We found that the optimal distance is 30 − 70 *μm*, corresponding to approximately 5 − 30 average number of neighbors per cell. This optimum may indicate the single cell local environment size that is most conserved, rather than broader structures within the tissue. Furthermore, even with a very low threshold of 10 *μm* where each node has only 0.02 neighbors per cell resulting in a graph with 98% of isolated nodes, or with a very high threshold of 90 *μm* where each cell has on average 50 neighbors, STELLAR performance does not degrade compared to using only molecular information. Overall, this analysis indicates that STELLAR is robust to input graph structure and that substantial performance gains can be expected by a meaningfully constructed spatial graph.

### STELLAR is applicable to spatial transcriptomics data

While existing methods are focused on either spatial proteomics or transcriptomics data, in addition to CODEX datasets we evaluated STELLAR on a single-cell transcriptome-imaging dataset generated using multiplexed error-robust fluorescence in situ hybridization (MERFISH). In particular, we applied STELLAR to a large-scale spatially resolved cell atlas of the mouse primary motor cortex [8] consisting of 23 granular cell types from two mice. We used a dataset from one mouse as the reference annotated dataset and evaluated performance on the dataset from another mouse. We then systematically removed a number of cell types from the reference set and evaluated how is STELLAR’s performance affected when gradually increasing the number of novel cell types. We measured accuracy separately on classes seen in the reference set and classes withheld from the reference set (novel classes). We find that STELLAR correctly assigned cell types to one of the classes in the reference set achieving 93% accuracy independently of the number of withheld cell types (Extended Data Fig. 7).

### STELLAR successfully annotates HuBMAP data

A critical current bottleneck in analysis of the spatially resolved single cells for consortia efforts is an accurate assignment of granular cell-type labels across donors of the same tissue. For example, a typical CODEX dataset with a 48-marker panel used to analyze four tissues of the intestine requires at least 25 hours of work to cluster, merge, recluster, subcluster, and assign cell types based on average marker expressions and locations within CODEX images. As a part of the HuBMAP consortium, we have generated CODEX imaging data for tissues from eight donors for a total of 64 tissues from a healthy intestine. Combined with single cell transcriptomic and epigenomic data, this molecularly detailed cell atlas contributes spatial organization of cell types as a reference map for understanding human intestinal biology [17].

We first evaluated the performance of STELLAR on this more granular data and used expert-annotated cell-type labels of images of different regions of a healthy colon generated from a single donor. We evaluated STELLAR in the leave-one-region-out setting where we trained on three regions and predicted annotations on the fourth region. STELLAR had high accuracy in cell-type label transfers (Fig. 3a) and had substantially higher performance (F-score of 0.8) than tested unsupervised clustering methods without manual intervention (F-score of 0.3 across different methods) [33]. This analysis showed STELLAR’s ability to recognize fine-grained level cell types and encouraged us to use STELLAR to annotate 2.6 millions of cells generated across 8 different donors.

**Figure 3:**
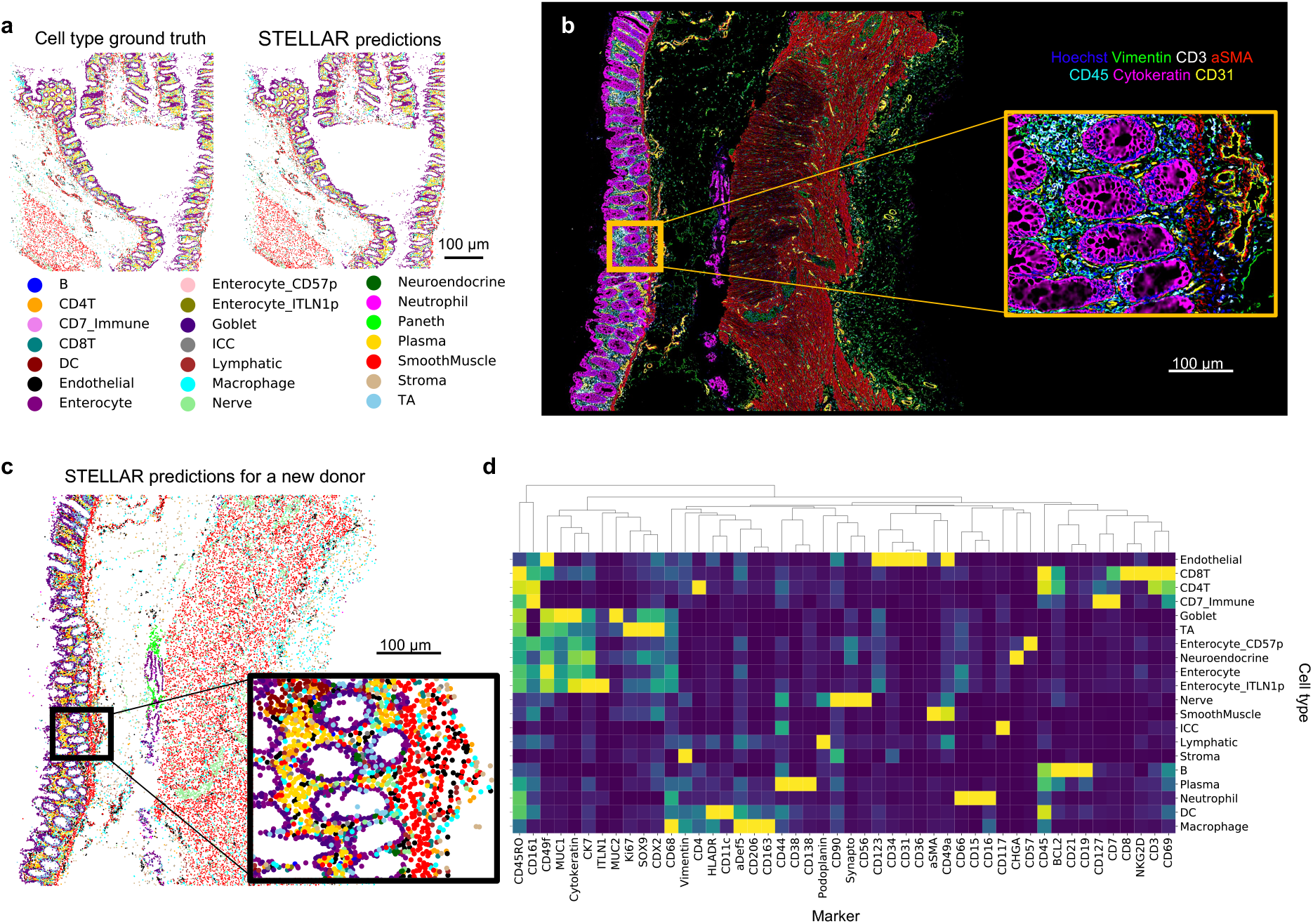
STELLAR transfers granular cell-type labels across tissue regions and donors from HuBMAP data and identifies major structures of healthy human intestine tissue. **(a)** Cell-type maps with ground-truth and STELLAR predictions made after training on CODEX multiplexed imaging of three different intestine regions across the colon. **(b-d)** We used data from one donor as the reference set, and applied STELLAR to annotate new, completely unannotated donor datasets. **(b)** Fluorescent images of 7 of 54 markers used in the CODEX multiplexed imaging of healthy intestine from a different donor. Hoechst, blue; vimentin, green; CD3, white; aSMA, red; CD45, cyan; cytokeratin, magenta; and CD31, yellow. Scale bar of the zoomed out image is 100 micrometers. **(c)** STELLAR predictions of cell types after training on data from a different donor mapped to CODEX spatial coordinates. **(d)** Average marker expression for cell types predicted by STELLAR for intestinal samples.

### STELLAR transfers information across donors

Encouraged by cross-region transfer results, we next used our expert-annotated samples from a single donor as training data and applied STELLAR to unannotated samples from two other donors. These datasets vary from each other based on tissue harvesting time, staining and imaging handler, and segmentation algorithms applied which may alter marker distribution. Despite the differences, we find that the fluorescent CODEX data of new donors (Fig. 3b and Supplementary Fig. 4) agree well with the cell-type map predicted by STELLAR and also agree with the known distribution of major cell types in the intestine (Fig. 3c and Extended Data Fig. 8). We additionally confirmed the quality of STELLAR’s predictions by looking at average marker expression profiles of predicted cell types for the samples from a different donor (Fig. 3d). Protein marker distributions match expert hand-annotated profiles in fine-grained cell types predicted by STELLAR. For example, CD4 is expressed only in cell types annotated as CD4^+^ T cells, whereas CD8 is expressed exclusively in CD8^+^ T cells. This additionally confirms that STELLAR predictions are reliable. We further applied STELLAR to six other donors, for a total of 8 donors, across 64 tissues and 2.6 million cells. STELLAR alleviated expert annotation effort tremendously – human annotations of these images would require approximately 320 hours of manual labor, while with STELLAR it took the expert only 4 hours to annotate the images.

We further validated STELLAR on datasets generated from different donors, that were prepared and stained at different times by different tissue handlers, were formulated into TMA (tissue microarray) or large tissue blocks, had different fixations conditions, used different image processing and cell segmentation algorithms, and used different antibody panels and clones for antibodies. This represents a particularly difficult case for cell type label transfer. We used fresh frozen CODEX intestine HuBMAP dataset as a reference dataset and applied STELLAR to a CODEX formalin-fixed, paraffin-embedded (FFPE) colorectal cancer dataset [25]. We used the subset of antigens targeted that overlapped even if the clone was different. While only a subset of the marker set overlapped, we find that STELLAR outperformed other methods by at least 22% (Supplementary Fig. 5).

### STELLAR embeddings capture higher-order tissue structures

The classification of cell-cell interactions enabled by multiplexed imaging of many cell types simultaneously unlocks the ability to identify and discover multicellular structures. Characterization of multicellular modules like tertiary lymphoid structures is critical to understanding tissue function, its relationship with disease, and how to design effective therapeutics [34]. We hypothesized that the latent cell embedding space learnt by STELLAR could reveal biologically meaningful information about the tissue organization. To explore this, we obtained cell embeddings from STELLAR and then clustered these cells in the STELLAR’s embedding space using the Louvain clustering [35] (Fig. 4a). We then evaluated the microenvironments represented by the resulting clusters by calculating the enrichment of cell types that neighbor these cells within these clusters as compared to tissue averages. This was also verified from looking at where the cell types with assigned cluster labels fell within the tissue using fluorescence staining and cell type neighbors to confirm larger pathological structures. We found that the clusters represent major multicellular structural features of the intestine such as the *immune follicle* structure, which was enriched for CD4^+^ T cells, B cells, and dendritic cells, as well as *secretory epithelial* which was enriched for goblet cells, neuroendocrine cells, and transit amplifying cells (Fig. 4b,c). We also confirmed this works across tissue sections taken at different sites of the colon with similar architectures (Extended Data Fig. 9). Similarly, we find that high-level structural information was detected also for the MERFISH data (Extended Data Fig. 10). These analyses revealed that the embedding space of STELLAR provides a method for identifying multicellular structures in tissues.

**Figure 4:**
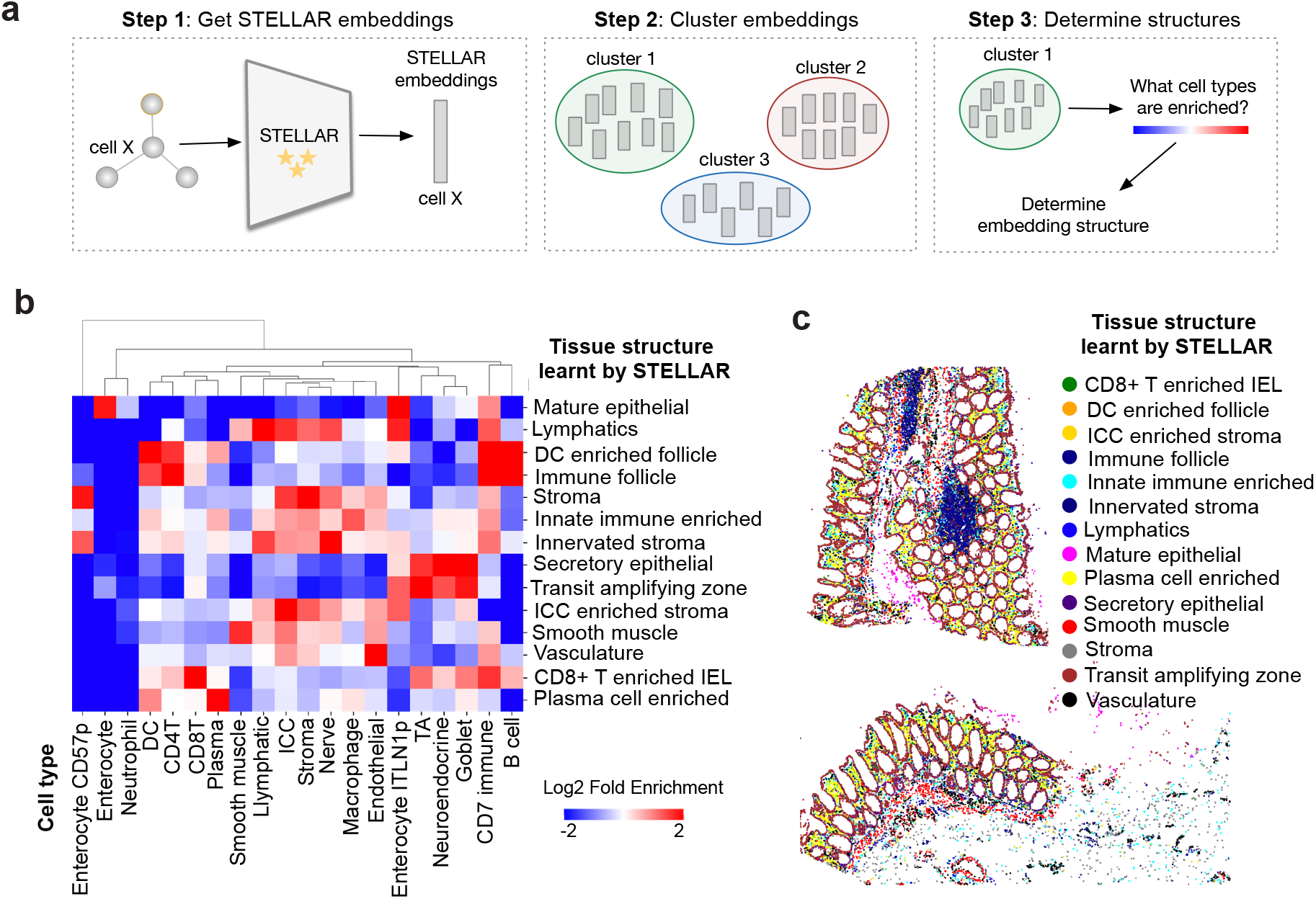
STELLAR’s embeddings reveal higher order tissue structures. **(a)** To analyze embedding space learnt by STELLAR, we clustered the cell embeddings using Louvain clustering. We then analyzed microenvironments of resultant clusters by checking which cell types are enriched in each cluster. The resultant clusters were manually annotated based on the cell type enrichments showed in (b). **(b)** Resultant clusters in STELLAR’s embedding space reveal that the embeddings capture major structures within the intestinal tissue. Heatmap shows enrichment of cell types assigned by STELLAR within the structural clusters as compared to tissue average percentages. **(c)** Graphical display of the tissue structures learnt by STELLAR found for one of the regions of the colon. Major structures such as immune follicles were identified by STELLAR.

## Discussion

STELLAR is an effective framework for annotating spatially resolved protein and RNA single-cell data and extracting spatial cell-interaction information. Two properties make STELLAR a unique tool in the single-cell toolbox: *(i)* the ability to learn low-dimensional cell embeddings that leverage spatial and molecular information across different biological contexts, and *(ii)* automatic identification of cell types from the reference set and discovery of novel cell types not present in the reference dataset. STELLAR learns a new cell embedding space that leverages spatial and molecular features using graph convolutional neural networks. The cell embeddings learnt by STELLAR can be used for any downstream analysis and provide a novel capability to identify major multi-cellular structures in the tissue. Our results support that incorporating spatial information directly leads to improved cell type annotation performance.

STELLAR is intended to be used for transferring annotations across datasets, including datasets from different biological contexts, *e.g.*, across dissection regions, donors, or related tissues. While doing so, STELLAR discovers expression patterns that define a novel cell type or cellular state. Our tool has a great value in transferring annotations across levels of granularity to a new biological context and discovering novel biological states that have not been characterized in previous experiments. The annotation transfer methodology in STELLAR distinguishes it from previous spatial annotation tools that rely on predefined marker genes that define cell types [16].

Currently, segmentation tools used largely rely on nuclear segmentation, and those that incorporate membrane based segmentation are far from perfect [36, 37]. Once segmentation masks and measurements become more reliable in regards to shapes of cells, then morphological features of cells could be included in STELLAR as feature vectors in addition to the marker expression. By including morphological features of cells, further improvements in performance could be expected as well as identification of unique cell states.

Development of STELLAR was motivated by a growing need to leverage spatial and molecular information across different biological contexts [17]. Multiplexed imaging technologies will drive future efforts to understand both healthy and diseased tissue processes, enabled by large consortia efforts that generate comprehensive datasets and standardize computational methods [5–7]. Novel insights will come from the rich, but still unexplored and underused information about spatial cell organization. We anticipate that STELLAR will alleviate future annotation efforts of large-scale spatial cell atlases, identify major multicellular structures in tissues, and reveal how cells cooperatively coordinate to enable tissues to function.

## Acknowledgements

This work was supported by the U.S. National Institutes of Health (2U19AI057229-16, 5P01HL10879707, 5R01GM10983604, 5R33CA18365403, 5U01AI101984-07, 5UH2AR06767604, 5R01CA19665703, 5U54CA20997103, 5F99CA212231-02, 1F32CA233203-01, 5U01AI140498-02, 1U54HG010426-01, 5U19AI100627-07, 1R01HL120724-01A1, R33CA183692, R01HL128173-04, 5P01AI131374-02, 5UG3DK114937-02, 1U19AI135976-01, IDIQ17X149, 1U2CCA233238-01, 1U2CCA233195-01); Cancer Research UK (C27165/A29073); the Parker Institute for Cancer Immunotherapy. JWH was supported by an NIH T32 Fellowship (T32CA196585) and an American Cancer Society - Roaring Fork Valley Postdoctoral Fellowship (PF-20-032-01-CSM). We also gratefully acknowledge the support of DARPA under Nos. HR00112190039 (TAMI), N660011924033 (MCS); ARO under Nos. W911NF-16-1-0342 (MURI), W911NF-16-1-0171 (DURIP); NSF under Nos. OAC-1835598 (CINES), OAC-1934578 (HDR), CCF-1918940 (Expeditions), IIS-2030477 (RAPID), NIH under No. R56LM013365; Stanford Data Science Initiative, Wu Tsai Neurosciences Institute, Amazon, JPMorgan Chase, Docomo, Hitachi, Juniper Networks, Intel, KDDI, and Toshiba.

## Author Contributions Statement

MB, KC, JWH and JL conceived the research. MB, KC, JWH and YT performed research and analyzed results. MB, KC and JL contributed new analytical tools and created the algorithmic framework. JWH, MPS, and GPN generated and analyzed the data. JL, GPN and MPS supervised the research. All authors participated in interpretation and wrote the manuscript.

## Competing Interests Statement

M.P.S. is cofounder and advisory board member of Personalis, Qbio, January AI, Mirvie, Filtricine, Fodsel, Protos. RTHM, Marble Therapeutics, and Crosshair Therapeutics. G.P.N. has equity in and is a scientific advisory board member of Akoya Biosciences, Inc.

## Methods

### Overview of STELLAR

STELLAR learns spatial and molecular cell similarities that are transferable across different biological contexts, such as different dissections regions, donors, or tissues. Across different contexts, STELLAR learns to automatically assign cells to cell types seen in the annotated reference set, or forms novel cell types if cells have unique properties that are not present in the reference dataset.

Specifically, STELLAR starts with an annotated reference cell graph 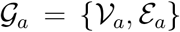 with molecular features for all nodes 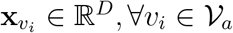, and an unannotated cell graph 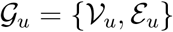 with molecular features 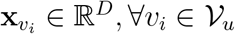. Here, 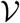 denotes the set of vertices and E denotes the set of edges in the graph 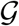. The nodes in each graph correspond to cells and cells are connected if they are spatially close. Node features correspond to gene or protein expressions of cells where *D* denotes the total number of measured genes/ proteins. For the reference graph we assume we are given a vector of cell annotations 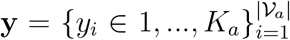 that assigns each cell to one of the *K_a_* cell types (or other annotations). STELLAR is ran on the shared subsets of the genes/proteins between annotated reference dataset and unannotated target dataset.

Given reference graph 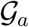 and unannotated graph 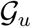, STELLAR first applies the encoder function *f_θ_* : ℝ^*D*^ → ℝ^*d*^ that maps cells from both graphs into a joint embedding space that captures spatial and molecular similarities between the cells. The cell embedding encoder function *f_θ_* is parameterized by learnable parameters *θ* of a graph convolutional neural network (GCN) [22]. The encoder function *f_θ_* generates *d*-dimensional cell embeddings 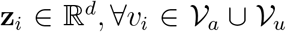. On top of the encoder function, we add a single linear layer parameterized by a weight matrix 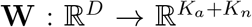, where *K_n_* corresponds to expected number of novel cell types that need to be discovered. Next, a softmax layer is added that assigns each cell to one of the *K_a_* + *K_n_* cell types.

### Graph construction

Given spatial cell coordinates of each cell, STELLAR first calculates Euclidean distances *d_i,j_* for each pair of cells (*v_i_*, *v_j_*) from the same region, and edge (*v_i_*, *v_j_*) is added to the edge set *ℇ* if *d_i,j_* < *τ*, where *τ* is a tunable threshold. We select the value of *τ* = 50*μm*. Graph construction step is independent of the subsequent method and can be changed as long as the constructed graph meaningfully reflects spatial similarities between cells.

### STELLAR encoder

The STELLAR encoder contains one fully connected layer followed by nonlinear activation function:

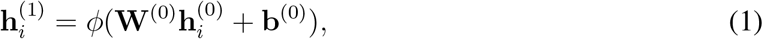

where 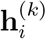 is the hidden state of node *v_i_* in *k*-th layer of the neural network and *k* = 0, 1. **W** is parameter matrix, **b** is bias vector and *φ* denotes nonlinear activation function. The hidden state 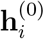 in layer 0 is set to node features 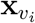 *i.e.*, gene or protein expression vector. The Rectified Linear Unit (ReLU) is used as the activation function *φ*: ReLU(·) = max(0, ·).

We then use a graph convolutional layer [22] to enable message passing among nearby cells:

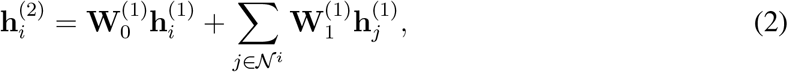

where 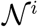 denotes neighborhood on node *v_i_*. The final embedding of node *v_i_* is 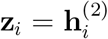. In non-spatial STELLAR (Fig. 2f), we replace the graph convolutional layer with another fully connected layer.

### STELLAR initialization

We initialize novel cell types with unlabeled data that lies out of distribution with respect to the labeled data. We first train a single-layer neural network *g_ψ_* on labeled data with cross-entropy loss. We cluster the unlabeled data using the Louvain algorithm with default parameters in scanpy [1] and then summarize the average entropy of each cluster with the trained *g_ψ_*. We select *K_n_* clusters with largest entropy and assign pseudo-labels corresponding to novel cell types as cells in the same cluster.

### STELLAR objective function

The objective function in STELLAR assigns cells to cell types from the reference set or discovers novel cell types. Inspired by [24], the objective function consists of two main components: *(i)* a component for discovering novel cell types, and *(ii)* a component for learning to recognize cell types from the reference set (Supplementary Fig. 1).

In the component for novel cell type discovery, we use an objective term that predicts pair-wise similarities given cell embeddings obtained using STELLAR encoder, *i.e.*, we predict whether two cells are similar or not. For the reference graph, we use ground-truth annotations to learn to predict similarity between two cells, that is two cells are similar if they belong to the same cell type. For the unannotated graph, pseudo-labels are generated based on the distances between cells in the embedding space. In particular, for each cell within the mini-batch, we identify the most similar nearest neighbor cell and generate pseudo-labels for the given pair. In that way, pseudo-labels are generated only for the pairs in which there is the most confidence. To find nearest neighbors, cell embeddings **h**^(1)^ without ReLU are used to allow cells from the reference graph to be selected as the neighbors of cells from the unannotated graph. Formally, the component for discovering cell types (DCT) minimizes the following term:

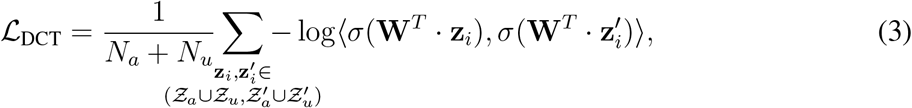

where **W** denotes linear layer weight matrix, *σ* denotes softmax function, *N_a_* and *N_u_* denote numbers of cells for annotated and unannotated graphs, respectively. 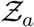 and 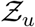 denote cell embeddings for annotated and unannotated graphs, and 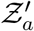 and 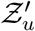 denote the set of closest neighbors of 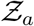 and 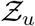 used for generating pseudo-labels.

The second component for recognizing cell types learns to distinguish cell types from the reference graph using ground-truth cell annotations **y**. Standard cross-entropy classification loss was enhanced with an adaptive margin mechanism that controls the learning speed of cell types from the reference set compared to novel cell types. Formally, STELLAR minimizes the following objective term to learn to recognize cell types (RCT) in the reference graph:

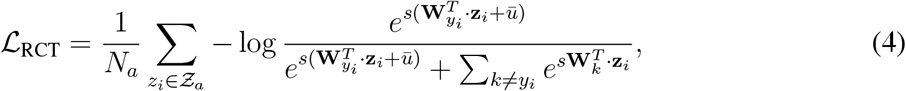

where *s* is temperature scaling parameter and 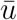 is uncertainty, 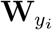 refers to the column vector that relates to class *y_i_*. The uncertainty is estimated as the average confidence of unlabeled examples computed from the output of the softmax function:

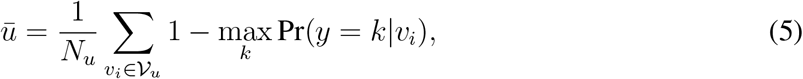

where *k* goes over all reference and novel cell types. At the start of the training uncertainty is large, leading to a larger margin and forcing larger intra-class variance [2]. As training proceeds, the margin becomes smaller and the objective boils down to standard cross-entropy.

Additionally, we use maximum entropy regularization term to avoid a trivial solution of assigning all cells to the same cell type. In particular, the regularization term is the following:

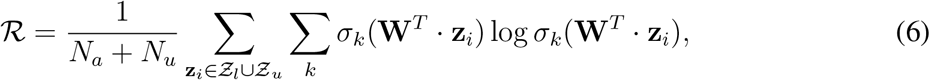

where *k* goes over all reference and novel cell types and *σ_k_* denotes *kth* cell type value of the softmax output.

Finally, the objective function in STELLAR combines reference cell type recognition, novel cell type discovery, and regularization components:

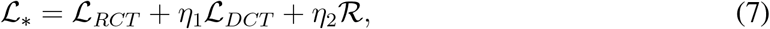

where *η*_1_ and *η*_2_ are regularization parameters.

### Architecture and hyperparameters

The encoder network in STELLAR consists of one fully-connected layer with ReLU activation and a graph convolutional layer with a hidden dimension of 128 in all layers. It uses the Adam optimizer with an initial learning rate of 10^−3^ and weight decay 0. The model is trained with a batch size of 512 for 20 epochs. A cluster sampler [3] first clusters input graphs into sub-graphs and then assigns the sub-graphs to mini-batches. The temperature scaling parameter *s* in eq. (4) is set to 10. Regularization parameters in (7), *η*_1_ add *η*_2_, are set to 1 and 0.3, respectively. These hyperparameters were used across all experiments.

### Number of novel cell types

STELLAR is initialized with the expected number of novel cell types as an input parameter. As the number of novel cell types is usually not known, STELLAR can be initialized with a large number of novel cell types and will automatically reduce the number by not assigning any cells to unneeded classification heads.

### Neighborhood identification analysis

Neighborhood analysis was performed as described previously [25]. Briefly, a window size of 10 nearest neighbors for each cell was taken across the tissue cell type maps. These vectors were overclustered to 20 clusters using k-means clustering algorithm. The clusters were mapped back to the tissue and evaluated for cell type enrichments to determine overall structure and merged down into final structures.

## Data Availability

The datasets presented in this study can be found in the online repository Dryad at https://datadryad.org/stash/share/1OQtxew0Unh3iAdP-ELew-ctwuPTBz6Oy8uuyxqliZk. Specifically, the quantified single-cell data are provided (with cells in rows and protein expression, x/y position, and cell type labels in columns). Additionally, we provide datasets used to transfer from the tonsil to BE tissue is there (BE Tonsil dryad.csv) and expert-annotated healthy human intestine (B004 training dryad.csv), which was used to test the accuracy of STELLAR across the four regions of the colon regions of this dataset and also for training for transferring cell-type labels to unlabeled donors (B0056 unannotated dryad.csv).

## Code Availability

STELLAR was written in Python 3.8 using the PyTorch library. The source code is available on Github at https://github.com/snap-stanford/stellar. The project website with links to data and code can be accessed at http://snap.stanford.edu/stellar/.

## Notes

### Competing Interest Statement

The authors have declared no competing interest.

### Summary of Updates

Figure updates

